# Implicit auditory memory in older listeners: from encoding to 6-month retention

**DOI:** 10.1101/2023.02.05.527176

**Authors:** Roberta Bianco, Edward T. R. Hall, Marcus. T. Pearce, Maria Chait

**Affiliations:** Ear Institute, University College London, WC1X 8EE London, United Kingdom; Neuroscience of Perception and Action Laboratory, Italian Institute of Technology, 00161 Rome, Italy; School of Electronic Engineering and Computer Science, Queen Mary University of London, E1 4NS London, United Kingdom; Department of Clinical Medicine, Aarhus University, 8000 Aarhus C, Denmark

**Keywords:** sequential pattern, sound perception, forgetting, statistical learning, Prediction by Partial Matching (PPM)

## Abstract

Any listening task, from sound recognition to sound-based communication, rests on auditory memory which is known to decline in healthy ageing. However, how this decline maps onto multiple components and stages of auditory memory remains poorly characterised. In an online unsupervised longitudinal study, we tested ageing effects on implicit auditory memory for rapid tone patterns. The test required participants (younger, aged 20-30, and older adults aged 60-70) to quickly respond to rapid regularly repeating patterns emerging from random sequences. Patterns were novel in most trials (REGn), but unbeknownst to the participants, a few distinct patterns reoccurred identically throughout the sessions (REGr). After correcting for processing speed, the response times (RT) to REGn should reflect the information held in echoic and short-term memory before detecting the pattern; long-term memory formation and retention should be reflected by the RT advantage (RTA) to REGr vs REGn which is expected to grow with exposure. Older participants were slower than younger adults in detecting REGn and exhibited a smaller RTA to REGr. Computational simulations using a model of auditory sequence memory indicated that these effects reflect age-related limitations both in early and long-term memory stages. In contrast to ageing-related accelerated forgetting of verbal material, here older adults maintained stable memory traces for REGr patterns up to 6 months after the first exposure. The results demonstrate that ageing is associated with reduced short-term memory and long-term memory formation for tone patterns, but not with forgetting, even over surprisingly long timescales.

## 1. Introduction

Memory loss is one of the most significant changes to cognitive processing experienced in healthy ageing (Cansino, 2009; Füllgrabe, 2020; Nyberg et al., 2012; Raz and Lindenberger, 2011; Salthouse, 2011). The most pronounced memory deficits are related to direct recall of episodic memory (Koen and Yonelinas, 2016, 2014), but evidence is increasingly revealing impairment in older listeners also in tasks that draw on automatic sensory memory processes (Cheng et al., 2013; Humes et al., 2013; Janacsek et al., 2012; Rieckmann and Bäckman, 2009; Rimmele et al., 2012; Schneider and Pichora-Fuller, 2000; Sluming et al., 2002; Wayne and Johnsrude, 2015).

Sensory memory is at the core of auditory processing (Atkinson and Shiffrin, 1968; Cowan, 1984; Nees, 2016; Winkler and Cowan, 2005). The nature of the unfolding signal is such that any listening task, from sound recognition to sound-based communication, depends on the ability to store successive events in memory to derive a coherent representation (Baldeweg, 2006; Conway, 2020; Heilbron and Chait, 2018; Massaro and Cohen, 1975; Rimmele et al., 2015; Santolin and Saffran, 2018). Age-related deficits in implicit auditory memory are increasingly being documented (Bianco and Chait, 2023; Fogerty et al., 2016; Jääskeläinen et al., 1999; Pekkonen et al., 1996; Rimmele et al., 2012), but remain poorly characterised due to limited computational tractability and paucity of longitudinal research designs. Recently emerging links between auditory processing and dementia are also making it urgent to quantify and understand these impairments (Griffiths et al., 2020; Johnson et al., 2021). In particular, the brain networks that are thought to underlie auditory memory – involving auditory, frontal and hippocampal areas (Barascud et al., 2016; Bonetti et al., 2022; Brown et al., 2004; Burunat et al., 2014; Kumar et al., 2014; Schapiro et al., 2014; Schmithorst, 2005; Watanabe et al., 2008) – are the same networks that exhibit age-related alterations (Mander et al., 2013) and the earliest decline in Alzheimer’s disease (Benhamou and Warren, 2020; Griffiths et al., 2020; Johnson et al., 2021). This makes auditory memory decline a promising proximity marker of dementia that is worth investigating (Fleischman, 2007; Swords et al., 2018).

Traditionally, auditory memory is conceived as a sequence of stores (Atkinson and Shiffrin, 1968): an echoic buffer stores detailed unprocessed information for several hundred milliseconds to allow successive sounds to be linked to the representation of sequences; then information passes to short-term memory for a few seconds and strengthens in long-term memory upon repeated presentations of the to-be-remembered sound (Winkler and Cowan, 2005). Impairments can potentially arise at any of these different stages.

Older, compared to younger, adults exhibit diminished amplitude and longer latencies of the mismatch negativity (MMN) – an automatic brain response evoked by a rare deviant sound in a sequence of standard sounds (Cooper et al., 2006; Jääskeläinen et al., 1999; Kiang et al., 2009; Pekkonen et al., 2001, 1996; Rimmele et al., 2012). The MMN is hypothesised to reflect the process of comparing incoming inputs with sensory-memory traces (Atienza et al., 2002; Baldeweg, 2006; Haenschel et al., 2005; Näätänen et al., 2007; Squires et al., 1976). Diminished MMN in older listeners suggests reduced echoic buffer and/or short-term memory (Näätänen et al., 2012). Age-related performance decline is also reported in probabilistic sequence learning tests using artificial auditory material and is hypothesised to reflect short- and long-term implicit memory deficits in older adults (Lukács and Kemény, 2015). In such tests, target sequences of arbitrary syllables or tones (presentation rate ∼2Hz) are structured according to certain probabilistic or deterministic statistics and repetitively presented to listeners unaware of the structure underlying the sequence (Christiansen, 2019; Petkov and ten Cate, 2020). Relative to novel sequences, a memory benefit for the target sequences is reflected by better immediate sequence reproduction or higher familiarity ratings after the exposure phase. This benefit declines with age and drastically at the age of around 65 (Herff et al., 2020; Lukács and Kemény, 2015; Schevenels et al., 2021).

Age-related decline has also been described in verbal memory tests asking participants to memorise a list of words, nonsense words, or stories to a minimum required level of accuracy. Memory is then typically probed with free recall at delayed sessions. Older compared with younger adults exhibit a reduced primacy effect suggesting age-related impairment in short-term memory (Mander et al., 2013; Murphy et al., 2000), as well as accelerated long-term forgetting (Elliott et al., 2014; Mary et al., 2013; McGibbon et al., 2022; Wearn et al., 2020) implicating impairments of long-term memory consolidation (Hoefeijzers et al., 2013). Notably, accelerated long-term forgetting has gained particular traction in the clinical field because of its potential as a predictor of Alzheimer’s pathology (Weston et al., 2018).

However, general confounding issues may emerge when tasks involve the presentation of stimuli at relatively slow rates (typically ∼ 2-4 Hz) and require engagement of the participant with the to-be-remembered information (e.g., the requirement to actively recall, judge familiarity etc). Factors such as experience, attention- or executive processing ability might conceal a core informational aspect of age-related memory decline. Indeed, age differences in immediate or long-term recall might reflect the effects of attentional load (Palmer et al., 2018), availability of feedback (Herff et al., 2020), or vocabulary knowledge (Schneider et al., 2002), as well as rehearsal strategies or interference (Manes et al., 2008; Mary et al., 2013; Wearn et al., 2020; Weston et al., 2018). Additionally, complex material, such as words, limits the cross-linguistic and translational diagnostic potential of verbal tests and, critically, it is difficult to informationally quantify and model. Consequently, whilst it is generally accepted that ageing is associated with auditory sensory memory impairment, the properties of this mnemonic decline are poorly understood. An additional persistent methodological hurdle lies in the ability to monitor the trajectory of memory processes over time – from the initial encoding of memories to their enduring retention in the long term. To achieve this, it is necessary to use standardized assessment techniques throughout these distinct stages.

We tested younger (aged between 20 and 30) and older (aged between 60 and 70) participants with an online paradigm that allows us to quantify short-term memory, the dynamics of long-term memory formation and long-term retention of tone patterns in an unsupervised manner. The “Auditory pattern Memory test” (ApMEM) (Bianco et al., 2020a) employs arbitrary rapid pure-tone sequences spanning the acoustic time scale of speech (20 Hz tone presentation rate, 1 Hz pattern rate) (Rosen, 1992). In 50% of the sequences, a pattern (REG; a repeating sequence of 20 tones) emerges partway, and listeners are required to detect it as quickly as possible (Fig. 1A). The rate at which successive tones are presented precludes deliberate tracking of the sequence structure, instead the REG patterns pop-out perceptually (Warren and Ackroff, 1976). Implicit auditory memory plays an obligatory role in pattern detection (Winkler et al., 2009). The brain’s ability to detect a REG pattern is hypothesised to arise from an automatic process that scans the unfolding sequence and compares incoming information with the just-heard sounds held in short-term memory and stored representations of the longer-term context (Barascud et al., 2016). Therefore, after adjusting for individual variability in detecting a simple sound change (see methods), the response time (RT) associated with REG detection becomes a quantitative measure of the information automatically held in memory until sufficient evidence has accumulated to trigger the observer’s behavioural response (Bianco et al., 2020a). In the task, most REG patterns are novel on each trial (REGn). The associated RT can thus be taken to reflect the combined contribution of implicit ***echoic and short-term memory*** to pattern detection. Crucially, unbeknownst to the participants, a few different patterns reoccur sparsely (REGr; every 2 minutes). Memory for REGr strengthens through repetitive exposure, as reflected by the gradual emergence of a RT advantage (RTA) in REGr pattern detection compared to REGn. This relative measure is used to quantify ***long-term memory formation***. Previous results from young adults have shown that this effect is implicit, in that it is not driven by explicit familiarity (Bianco et al., 2020a). To measure ***long-term memory retention*** of REGr patterns, RT to REGr is also measured 8 days and 6 months after the first exposure.

**Figure 1.**
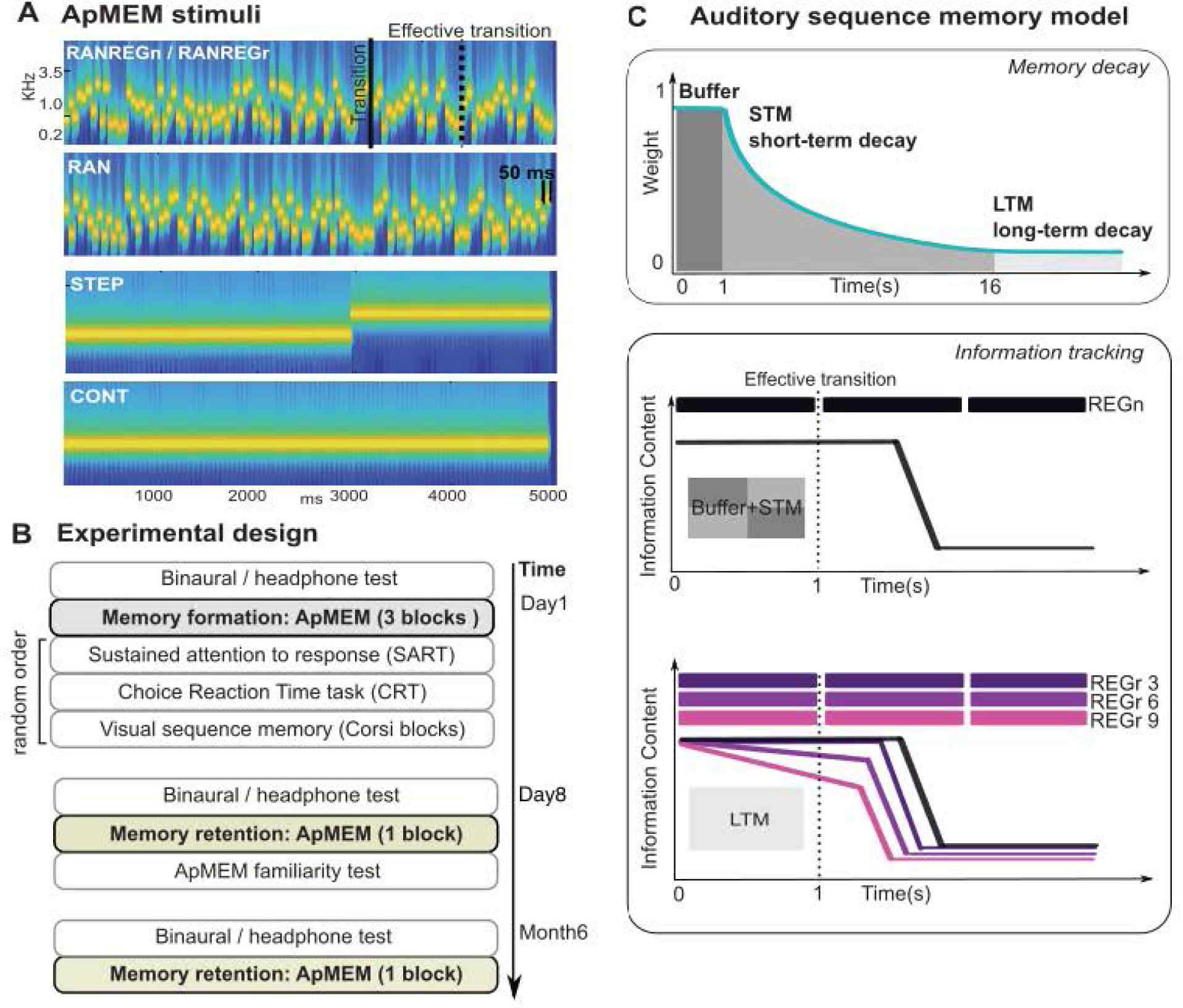
Stimuli, experimental design, and a model of auditory memory. **A) Example spectrograms of the ApMEM stimuli**. RAN sequences contained a random arrangement of tone pips. RANREG sequences contained a transition from a random (RAN) to regularly repeating cycles of 20 tone-pips (REGn). The repetition of the REG pattern becomes detectable after the first cycle (‘effective transition’). RAN and REGn sequences were generated anew on each trial. Three different regular patterns (REGr) were each presented identically trice within a block. Reoccurrences were spaced ∼2 min apart. STEP stimuli, containing a step change in frequency, (and their ‘no change’ control, CONT) were also included in the stimulus set as a form of attentional checks, and to control for age-related differences in speed of processing reflected in RT-based measures. To distil the computation time required to detect the patterns RTs to RANREG were corrected by the median STEP RTs. **B) Experiment design and task order.** This was a 3-session study, conducted on day 1 (d1) to test memory formation over 3 blocks of ApMEM task, and on day 8 (d8) and month 6 (m6) to test memory retention with 1 block of ApMEM. Additional cognitive tests (random order) were included on d1: Sustained attention to response (SART), Choice reaction time task (CRT), and visual sequence memory task (Corsi blocks). The ApMEM familiarity surprise task was administered at the end of d8. **C) Auditory sequence memory model.** A schematic representation of the parameters implemented in the model to simulate auditory memory. **Memory decay.** Sequence statistics are memorised through partitioning the unfolding sequence into events and sub-sequences of increasing order (n-grams) that are thereon stored in memory. The salience of these observations (‘weight’) decays over time through 3 phases indicated by the shaded areas: (1) buffer, (2) short-term memory (STM, with a fast exponential decay from the weight of the buffer to the starting weight of the next phase over a given duration), and (3) the long-term memory (LTM, with a slow exponential-decay defined by a starting weight and a half-life). **Information tracking:** The model uses these stored statistics to quantify the information content (IC, where high IC corresponds to low predictability and low IC corresponds to high predictability) of incoming tones. A newly encountered pattern (REGn, in black) is detected when incoming information matches the information held in the buffer and the STM resulting in a drop of IC. Once an n-gram is encountered, its representation enters a slow LTM-decay phase. Pattern reoccurrence (REGr, illustrated with different shades of purple to indicate the same pattern after 3, 6 and 9 reoccurrences), leads to weight increase in memory as an index of increasing memory strength. This causes the fall in IC associated with pattern recognition to occur progressively earlier. Previous observations (Barascud et al., 2016; Bianco et al., 2020a) support the hypothesis that listeners track and leverage the drop in IC as an indication of the emergence of regularity.

Utilizing such arbitrary stimuli presents several notable advantages when investigating auditory memory. Beyond overcoming potential linguistic barriers, these stimuli are too rapid to be actively monitored (Warren and Ackroff, 1976), and unlikely to be encountered in real-world settings. Consequently, they effectively mitigate mnemonic biases stemming from prior linguistic familiarity, deliberate rehearsal tactics, or the interference posed by routinely encountered auditory inputs. Importantly, these stimuli are also well suited for establishing a systematic link between implicit listener performance (measured through RTs to patterns) and the information-theoretic attributes of the stimuli. This facilitates the computational simulation and quantification of how mnemonic subcomponents constrain listener behaviour.

The Prediction by Partial Matching (PPM) model, has successfully explained listeners’ performance with a variety of discrete musical and artificial auditory sequences (Barascud et al., 2016; Bianco et al., 2020b; Di Liberto et al., 2020; Harrison et al., 2020; Kern et al., 2022; Omigie et al., 2019; Pearce et al., 2010; Quiroga-Martinez et al., 2020). Here, we use its memory-constrained variant (PPM-decay, see (Harrison et al., 2020), previously used to simulate performance in the ApMEM task in young listeners’ (Bianco et al., 2020a). The model encodes sequences by weighting sub-sequences (n-grams) of multiple orders based on recency. Each auditory event is stored as a single count with a certain weight which decays over time (following the parameters set by a customizable decay kernel; Fig. 1C) and increases if the event is re-encountered. The non-linear decay profile simulates the contribution of three subcomponents to memory formation corresponding to echoic buffer, short- and long-term memory decay. Based on stored observations, the model estimates the information content (IC) for each event, corresponding to the degree to which the event is expected based on prior context. When a random sequence transitions into a novel REG pattern (REGn), the IC drops due to the match between incoming information and prior observations held in short-term memory. The brain is hypothesized to be sensitive to this change in IC (Barascud et al., 2016) as an indicator for changes in underlying stimulus statistics. For a previously encountered REG pattern (REGr), there will be a stronger match with information stored in long-term memory, resulting in earlier change in IC and consequently a faster response (Fig 1C). Within this framework, the RT to a pattern may encapsulate a combination of both speed in accessing stored information and associated confidence indirectly reflecting the weights this information has in memory.

Fitting ApMEM RT data from young and older adults with parameters associated with the decay kernel and memory weights, allowed us to identify the potential sources of memory impairment in the ageing cohort.

## 2. Methods

### 2.1 Power analysis

We initially ran an online pilot experiment (N = 20, age between 20-30 years old). The RTA effects size across 3 blocks was h_p_^2^ = .22. We expected the difference between groups to be potentially small (h_p_^2^ = .02). A prospective power calculation (beta = 0.8; alpha = .05) for an ANOVA within-between interaction yielded a required total sample size of N = 41 per group. We set our online target sample size to N = 90 per group to account for dropouts (expected ∼30%) due to headphone check exclusion and the unsupervised and longitudinal nature of the experiment. Experimental procedures were approved by the research ethics committee of University College London and informed consent was obtained from each participant.

### 2.2 Participants

Two participant groups were recruited via the Prolific platform (https://www.prolific.co/). A group of younger participants (age range 20-30 years old; N=93) and a group of older participants (age range 60-70 years old; N=98). Inclusion criteria included being a native speaker of British English, generally good health, no known hearing problems, or cognitive impairment (all based on self-report). Participants using low-quality audio equipment, or those suffering from binaural hearing loss, were screened out using the test introduced by Milne et al. (2020). 29 participants in the older group, and 24 of the younger participants failed the screen and their data were therefore not analysed. Additional exclusion criteria were: (a) poor performance in the main apMEM test (mean d’ < 1.5 across blocks in day 1 or day 8; N = 1 in the older group, N = 4 in the younger group excluded.) (b) poor performance on the attentional checks (mean RT to STEP changes larger than 2 STD away from the group means; N=1 in the younger group excluded). A final N = 132 was analysed: N = 68 (28 female) in the older group, and N = 64 (33 female) in the younger group (Fig. 2A). Six months later we ran an additional (surprise) session. From the original pool, N = 104 participants (N = 63 older, N = 41 younger group) signed up, and, after exclusion as mentioned above, data from 93 subjects were analysed.

**Figure 2.**
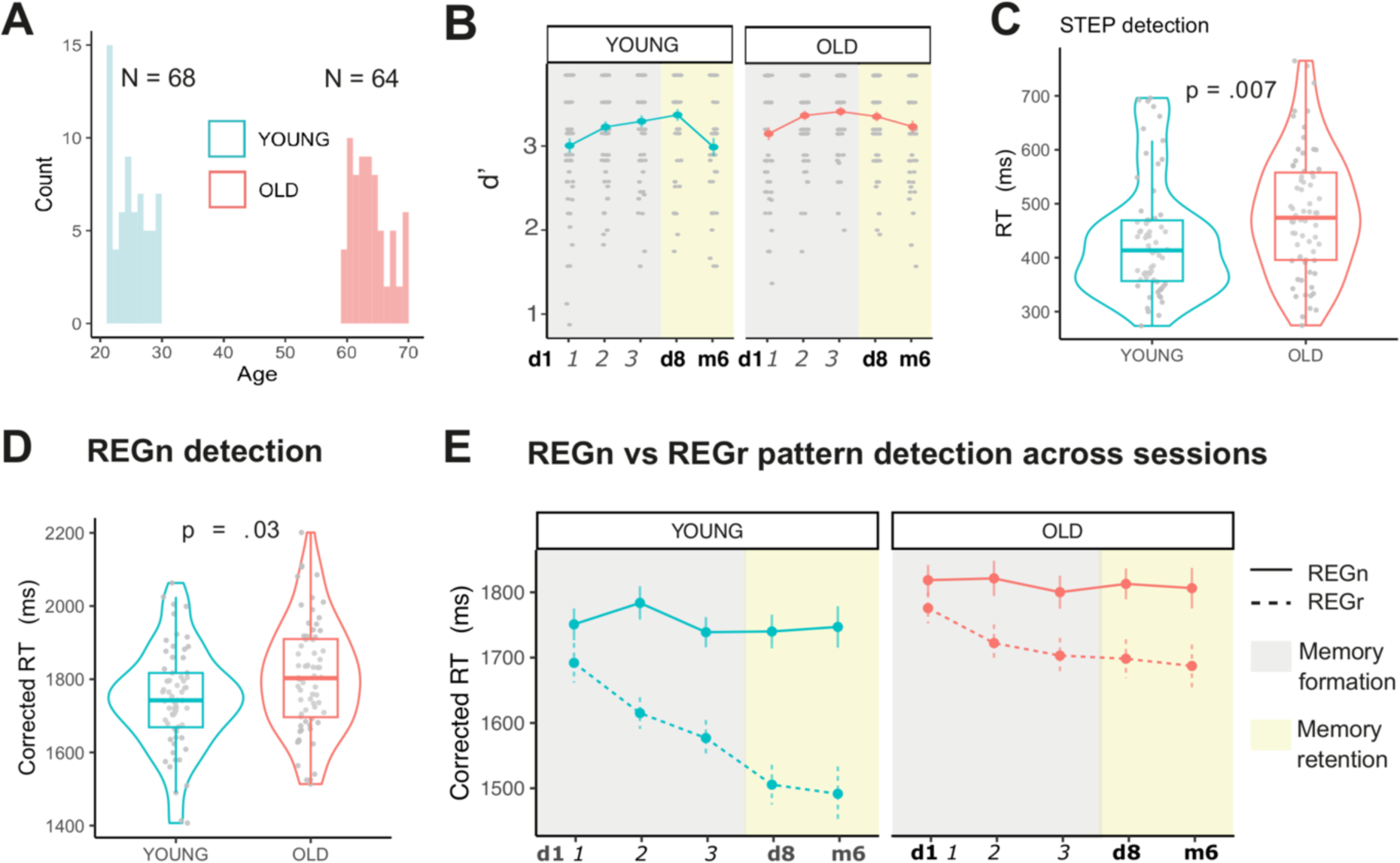
Auditory memory formation in young and older adults (ApMEM task). **A)** Participant age distribution. **B)** d’ (sensitivity to the emergence of regularity) for the OLD and YOUNG groups. Shaded areas indicate the different stages of memory formation (d1 in grey) and retention (d8, m6 in yellow). **C)** Median RT to STEP trials (frequency step changes) averaged across the d1, d8 and m6 sessions. **(D)** Across all sessions (d1, d8 and m6), older listeners exhibited slower REGn RT than young controls. **E)** RTs to REGr and REGn across the 3 blocks of d1, and the 1 block of d8 and m6. Error bars represent the standard error of the mean.

### 2.3 General procedure

This study was implemented in the Gorilla Experiment Builder platform (www.gorilla.sc) (Anwyl-Irvine et al., 2020) and delivered across three sessions: day 1 (d1), day 8 (d8) and month 6 (m6) (Fig. 1B). Participants were initially recruited only for d1 and d8. They were later invited to participate in the m6 session. Participants were recruited via the Prolific platform and remunerated based on an hourly wage of £ 8. To enhance the precision of online RT measurements and minimize the potential interference arising from variances in devices and browsers (as outlined by Anwyl-Irvine et al., 2020; Bridges et al., 2020), we implemented a controlled approach. Only participants equipped with a computer (excluding tablets or phones) and using the ‘Chrome’ browser were eligible to take part in the study. Offline checks of the OS used by participants revealed no systematic bias between older and younger participants (see Table S1 in supplementary material). Potential biases in laptop vs desktop computers or OS versions were not assessed. Participants who performed below 70% accuracy in the practice of the main ApMEM task (see below) were prevented from continuing and received partial compensation for the time spent on the experiment.

On **day 1 (60 minutes)**, participants first completed a headphone check (Milne et al 2020; strict test version). The test is based on a binaural pitch signal that is only audible over headphones (i.e., where L and R audio channels are delivered separately to each ear). Passing the test requires reasonable quality audio equipment (headphones with separate R and L channels) and preserved binaural hearing (Sanchez Lopez et al., 2018). People who failed this very first stage were excluded from the analysis. Next, participants performed the Auditory pattern Memory task (ApMEM; 3 blocks). The main task was preceded by a short practice with a simplified version of the stimuli (see below). People who did not reach 70% accuracy in this practice stage were stopped from continuing the experiment and received partial compensation. ApMEM was followed by a series of cognitive tests - Sustained Attention to Response Task (SART), the Corsi blocks task, and the Choice Reaction Time (CRT) task - presented in random order across participants (Fig. 6A). More details about each task are provided below. At the end of the session, participants completed a short questionnaire about their listening environment and equipment. This included 4 choice questions about how old (1 year/2-4 years/5-7 years/8 or more), and how expensive their computer was (less than £700 / between £700 and £1000 / between £1000 and £2000 / more than £2000). We observed no between-group differences associated with these two ratings (W = 2100, p-value = 0.711; W = 1855, p-value = 0.100). Participants were also asked about their physical activity habits (numbers of hours per week) and years of musical training (ranked as 0, 0.5, 1, 2, 3-5, 6-9, 10 or more).

On **day 8 (15 minutes)**, participants completed the headphone check followed by a single ApMEM block. The session ended with a surprise familiarity test for REGr (see details below). **6 Months** later participants who completed d1 and d8 were re-invited for another surprise session **(15 minutes)**. This included the headphone check and 1 block of ApMEM.

### 2.4 Tasks

#### 2.4.1 Headphone check

This test was used to exclude from the analysis participants with poor sound environments. We used the strict version of the test freely available online (https://gorilla.sc/ openmaterials/100917) and thoroughly documented in (Bianco et al., 2021; Milne et al., 2020).

#### 2.4.2 ApMEM task

The ApMEM task was used to measure multiple stages of auditory memory (Bianco et al., 2020a). Stimuli (Fig. 1A) were sequences of contiguous 50-ms tone-pips of different frequencies generated at a sampling rate of 22.05 kHz and gated on and off with 5-ms raised cosine ramps. Twenty frequencies (logarithmically spaced values between 222 and 2,000 Hz; 12% steps; loudness normalised based on iso226) were arranged in sequences with a total duration varying between 5.5 and 6 s. The specific order in which these frequencies were successively distributed defined different conditions that were otherwise identical in their spectral and timing profiles. We created 5 different stimulus sets which participants were randomly assigned to. Each set contained five conditions as follows. RAN (‘random’) sequences consisted of tone-pips arranged in random order. This was implemented by sampling uniformly from the pool with the constraint that adjacent tones were not of the same frequency. Each frequency was equiprobable across the sequence duration. The RANREG (random-to-regular) sequences contained a transition between a random (RAN), and a regularly repeating pattern: Sequences with initially randomly ordered tones changed into regularly repeating cycles of 20 tones (an overall cycle duration of 1 s; new on each trial). The change occurred between 2.5 and 3 s (random jitter) after sequence onset such that each RANREG sequence contained 3 REG cycles. RAN and RANREGn (RANREG novel) conditions were generated anew for each trial and occurred equiprobably. Additionally, and unbeknownst to participants, 3 different REG patterns reoccurred identically several times within the d1, d8 and m6 sessions (RANREGr condition, reoccurring). The RAN portion of RANREGr trials was always novel. Each of the 3 regular patterns (REGr) reoccurred 3 times per block (every ∼ 2 minutes, i.e., 9 presentations in d1, 3 in d8, and 3 in m6). Reoccurrences were distributed within each block such that they occurred at the beginning (first third), middle and end of each block. Two control conditions were also included: sequences of tones of a fixed frequency (CONT), and sequences with a step change in frequency partway through the trial (STEP). The STEP trials served as a lower-bound measure of individuals’ RT to simple acoustic changes. They were also used as attention checks – no, or very slow (see below) responses to STEP trials indicated insufficient task engagement.

Each session of the main task was preceded by a volume adjustment stage. Participants heard a few sounds from the main task and were instructed to adjust the volume to a comfortable listening level. In the main task, participants were instructed to monitor for transitions (50% of trials) from random to regular patterns (RANREG) and frequency changes in STEP stimuli and press a keyboard button as soon as possible upon pattern detection. On day 1, to acquaint participants with the task, two practice runs were administered. The first practice contained 24 sequences consisting of simplified versions of the stimuli (10 RAN, 10 RANREGn, 2 STEP, 2 CONT), in that sequences were presented at a slower tempo (10 Hz) and contained regularities of 10 tones. The second practice consisted of 21 sequences (9 RAN, 9 RANREGn, 2 STEP, 1 CONT) presented at a faster tempo (20 Hz) and containing regularities of 20 tones, as in the main task. The main task consisted of 3 blocks on d1, 1 on d8 and 1 on m6 sessions. Each block lasted about 6 minutes and contained 43 stimuli (18 RAN, 9 RANREGn, 9 RANREGr, 5 STEP, 2 CONT), with an inter-trial-interval of 1 s. Participants were instructed to respond as quickly and accurately as possible both to the transition from random to regular pattern and to the step frequency change. Feedback on accuracy and speed was provided at the end of each trial as in our previous work (Bianco et al., 2020a): a red cross for incorrect responses, and a tick after correct responses. The colour of the tick was green if responses were ‘fast’ (< 2200 ms from transition to REG or <500 ms from the step frequency change), and orange otherwise. The inter-block intervals were set to have a maximum duration of 3 minutes to keep the overall duration of the exposure equal across participants. Altogether, on day 1 instructions and practice took approximately 20 min and the main task lasted 18 minutes. On day 8 and month 6, the ApMEM task took 8 min, 2 of which consisted of 20 trials of practice.

d’ (computed per session across RANREGn and RANREGr conditions) served as a general measure of sensitivity to regularity. Responses were marked as hits when they occurred after the pattern begins to repeat (i.e., after the first cycle; effective transition). Responses that occurred earlier or during random trials were marked as false alarms. Participants whose d’ was smaller than 1.5 in at least one session were excluded from the analysis as this indicated poor pattern sensitivity and made RT uninterpretable. The core analysis focused on the RTs to the onset of regular patterns and followed the pipeline adopted in (Barascud et al., 2016; Bianco et al., 2020a). RT was defined as the time difference between the onset of the regular pattern or the frequency step change and the participant’s button press. For each participant and block, RTs beyond ±2 SD from the mean were discarded (OLD: 4.3% in REG, 4.1 in REGr; YOUNG: 4.1% in REG, 3.8% in REGr). Individuals identified as outliers in the RTs to the STEP condition were excluded from the analysis as this indicated low task engagement. The median STEP RT computed per session was used as a measure of the latency of the response to a simple acoustic change. This measure of individual baseline was stable across sessions (day 1 vs day 8 OLD: V = 991, p = .267; YOUNG: V = 1012, p = .854), and was subtracted from the RTs to RANREGn and RANREGr to yield a lower-bound estimate of the computation time required for pattern detection. Lastly, for each subject, we computed indexes of RT advantage (RTA) of REGr over REGn to quantify memory at different time points. To do so, we first corrected the RTs to REGr trials by the median RTs to REGn in each block. Then, to calculate the RTA by block, we computed the median RTA across the 3 intra-block presentations and the 3 different REGr patterns.

#### 2.4.3 ApMEM familiarity task

Explicit memory for REGr was examined with a surprise task at the end of day 8. The 3 REGr patterns presented in the ApMEM (only one instance per REGr) were intermixed with 18 REGn patterns, as in (Bianco et al., 2020a). Participants were instructed to indicate which patterns sounded ‘familiar’. The task took approximately 2 minutes to complete. The classification was evaluated using the Matthews Correlation Coefficient (MCC) score which ranges between 1 (perfect classification) to −1 (total misclassification) (Boughorbel et al., 2017; Powers, 2007). Before starting the task, participants were played a few sounds similar to those played in the upcoming task and asked to adjust the volume to a comfortable listening level.

#### 2.4.4 Choice reaction time task (CRT)

The CRT task is an established measure of individual variability in processing speed and is known to be linked with age-related decline in higher-level cognitive functions (Salthouse, 1996). Subjects were required to respond as soon as possible with the index or middle finger to a cue appearing with equal probability on the left or right box displayed on the screen. The task comprised 20 trials and took approximately 1 minute to complete. The task has two outcome measures: The central tendency (the median RT), and intraindividual variability (the raw standard deviation of the RTs), known to show a marked increase with age (Der and Deary, 2006; Hultsch et al., 2002).

#### 2.4.5 Corsi blocks (visual-sequence memory) task

Nine identical black squares were presented on the screen. On each trial, following a fixation duration (500 ms) a few blocks flashed (briefly changed colour from black to yellow; flash duration 500ms; inter-flash-interval 250 ms) in a sequence. Participants had to reproduce the order of the sequence by mouse-clicking on the correct blocks. The initial sequence length was 2 blocks. Correct responses resulted in a length increase and incorrect responses in a length decrease. Overall participants completed 20 trials. The task took approximately 5 minutes to complete. As an outcome measure, we computed the mean sequence length. This score is considered to reflect the ability to remember the temporal order of spatial sequences and it is known to deteriorate with ageing (Beigneux et al., 2007; Bianco and Chait, 2022; Fournet et al., 2012).

#### 2.4.6 Sustained Attention to Response test (SART)

The ApMEM task is attentionally demanding and memory formation may be affected by the listener’s capacity to sustain focused attention. The SART task was used to measure individual vigilance and propensity to inattention (Manly et al., 2000). Participants were asked to respond by pressing a button to serially and frequently presented ‘go’ visual stimuli (digits from 0 to 9, except 3) but maintain a readiness to withhold a response to rare and unpredictable no-go trials (the digit 3). The task took approximately 8 minutes to complete. The key outcome measure was the % ‘no-go’ fail – quantifying listeners’ ability to successfully stay “on task”.

### 2.6 Statistical analyses

Performance was statistically tested with linear analyses of variance (ANOVA) implemented in the R environment using the ‘ezANOVA’ function (Michael Lawrence, 2016). P-values were Greenhouse-Geisser adjusted when sphericity assumptions were violated. Post hoc t-tests were used to test for differences in performance between conditions across blocks and groups. A Bonferroni correction was applied by multiplying p values by the number of comparisons. Resulting values below the significance level of .05 are indicated as n.s. – non-significant. Non-parametric tests were used where the normality of the outcome distribution and homogeneity of variances were violated based on the Shapiro-Wilk test. To isolate the contributions of different tasks to ApMEM performance, we used hierarchical linear regressions.

### 2.6 PPM-decay modelling

Observed data from the ApMEM task were computationally modelled using a memory-constrained Prediction by Partial Matching (PPM) model. PPM is a variable-order Markov modelling technique that estimates the likelihood of the occurrence of symbolic sequential events, given the number of occurrences of n-grams of varying size within a training sequence, smoothing between models of different orders (Bunton, 1996; Cleary and Witten, 1984).

Conventional models using PPM possess a perfect memory for all events in their training data, regardless of proximity to the modelled event. To model the effects of human memory on learning, Harrison et al. (2020) implemented a PPM model with the ability to down-weight occurrences in the model over time, based on a customisable decay kernel. As used here, the kernel contained three phases: (1) a high-fidelity echoic memory buffer, defined by a weight and a duration; (2) a short-term memory (STM) phase that decays exponentially from the weight of the buffer to the starting weight of the next phase over a given duration; and (3) an exponentially decaying long-term memory (LTM) phase, defined by a starting weight and a half-life; (examples of decay kernels and their phases can be seen in Fig. 3C). Additionally, varying levels of noise were added to event probabilities, fulfilling the role of a general processing speed parameter, and replicating similar imperfections in human memory. This noise was parameterised by the SD of a normal distribution, from which sampled absolute values were added to the weights on memory retrieval.

**Figure 3.**
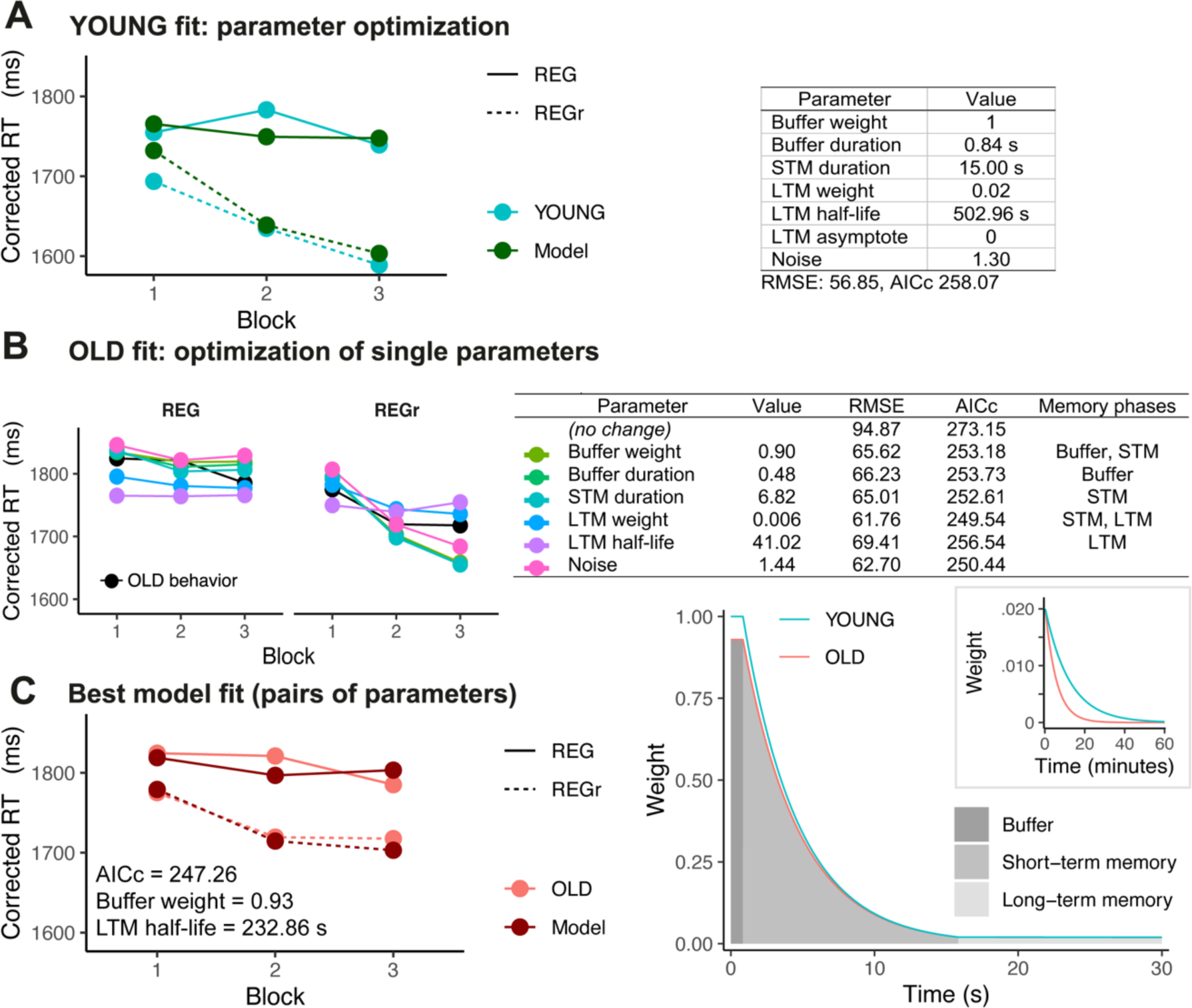
Modelling contribution of buffer, STM and LTM to auditory memory formation in young and older adults. **A)** Simulated and observed RTs to REGn and REGr conditions for the YOUNG group (left), and the parameter values of the optimised model with RMSE = 56.85 (right). **B)** Simulated RTs and parameter values for models fitting OLD RTs by optimising individual parameter changes from those of YOUNG. **C)** Simulated and observed RTs to REGn and REGr conditions for the OLD group using parameter values obtained for YOUNG, modifying buffer weight and LTM half-life yielding RMSE = 57.22 (left), and the memory decay kernels of best fitting models for both groups on a time scale of 30 seconds (right). The small insert illustrates the LTM phase on a timescale of 1 hour. See Supplementary Material for simulated RTs using tone intervals, instead of absolute tone values.

All stimuli in blocks 1 to 3 of day 1 were modelled, as presented for each stimulus set, maintaining the tone, stimulus, and block timings of the task. Models were trained dynamically, estimating a probability for each tone, given the sequence preceding it and all preceding stimuli, which was converted into information content (negative log-base-2 probability). Models were limited to a maximum n-gram length of 5 symbols (an order bound of 4). As in (Bianco et al., 2020a), changes in information content were identified for REGn and REGr stimuli using the nonparametric change-point detection algorithm of (Ross et al., 2011), a sequential application of the Mann-Whitney test, while controlling for a Type I error rate of 1 in 10,000.

Model parameters were optimised to find the decay configuration that best reproduced the observed data of the younger group using Rowan’s Subplex algorithm, as implemented in the NLopt package (Johnson, 2020; Rowan, 1990). Initial parameter values were adapted from the manually fitted parameters by Bianco et al. (2020a). To account for the increased variability of change points due to modelling prediction noise, for every optimisation iteration, the modelling was repeated 100 times, refreshing model memory between each. Repeated change points were then averaged for individual stimuli. Optimisation sought to minimise the root-mean-square error (RMSE) between observed RTs and modelled change points, when averaged for each of the REGn and REGr conditions within blocks 1 to 3, for each different stimulus set. To facilitate a comparison of goodness of fit between models using different numbers of parameters to fit the same data, Akaike Information Criterion (AIC) values were calculated under the assumption of normally distributed errors. A correction was applied to penalize potential overfitting when using small sample sizes (AICc).

To characterise differences between the older and younger groups, first, observed data for the older group were modelled by optimising a single parameter while holding all others to the values obtained for the younger group. The fit of these models is shown in Fig. 3B. As no single change of only an individual parameter adequately reproduced the observed data, pairs of parameters were then fitted. For each pair, the parameter giving the best fit when optimised individually was initialised at the value obtained in that optimisation. The other parameter in the pair was initialised at, and all remaining parameters were held at, the values obtained for the younger group. The parameters that produced the best-fitting of these models, based on RMSE, were selected as those characterising the older group.

## 3. Results

With an online version of ApMEM, we characterised implicit memory for rapid tone patterns in young and older participants over multiple timescales, from early mnemonic stages required to detect novel patterns to long-term memory formation and retention at 1-week (d8) and 6 months (m6). Using a computational model of auditory sequence processing, we then distilled and quantified how different memory parameters contribute to between-group differences in performance on day 1 (d1).

To account for age-related effects on general aspects of the ApMEM task (ApMEM task is RT-based, measures memory for sequences and requires focused attention) we also included tests of processing speed (Salthouse, 1996), spatial-visual sequence memory (Corsi, 1972; Kessels et al., 2000), and attention (Manly et al., 2000) (Fig. 1B). We explored whether these measures might explain response differences in ApMEM performance between the age groups.

### 3.1 No group difference in accuracy of pattern detection

Overall, the ability to detect the emergence of a pattern from a random sequence, as quantified with d’ (collapsed across REGn and REGr), was consistently high across blocks (Fig. 2B), though a drop between d8 and m6 was seen in the younger group (YOUNG: W = 796, p = 0.003, CI [−0.642 −4.953e-05; OLD: W = 1628, p = 0.208, CI [−0.321 3.74e-05]). d’ was similar between groups on d1 (W = 2524.5, p = .113, CI [−.006 .231], mean OLD: 3.3 ± .40, YOUNG: 3.17 ± .454), on d8 (W = 2036, p =.513, CI [−0.321 7.035e-06], mean OLD: 3.34 ± .48, YOUNG: 3.36 ± .558), and on m6 (W = 1265, p = 0.082, CI [−1.999e-05 .496], mean OLD: 3.22 ± .55, YOUNG: 2.97 ± .67). This confirms high sensitivity to the presence of regularities and allows us to confidently interpret the between-group differences in RTs as a measure of memory strength.

The hit rate for novel (REGn) and reoccurring patterns (REGr) was computed for each session (d1, d8, m6). Hits were higher for REGr than REGn on d1 (mean over blocks 1,2 and 3) in both groups (REGr: 96.6±4.36, REGn: 94.1±7.7 in OLD; REGr: 97±4.57, REGn: 93.6±7.9 in YOUNG; one-sample Wilcoxon test of REGr – REGn hit per cent, OLD: V = 596, p = .006; YOUNG: V = 807, p < .001), indicating a memory effect for REGr. The hit advantage of REGr over REGn did not differ between the groups on d1 (W = 2439.5, p = .218, CI [−2.86e-05 3.70]), on d8 (W = 2341.5, p= .39, CI [−1.38e-05 3.97e-05]), and m6 (W = 1118.5, p = .531, CI [−3.47e-05 3.44e-05]). Finally, the hit advantage did not change over time: similar effects were found in the last block of d1 (b3_d1), d8 and m6 in both groups (all p < .78).

Overall, both groups were highly accurate in detecting the patterns and showed a memory advantage for REGr. However, this advantage was similar between groups and did not change over time, likely because it quickly reached ceiling effects. The ApMEM task was designed to yield high levels of accuracy consistently over time. This enabled us to focus on analysing the RT to capture processes related to dynamic sequence processing. Below we demonstrate that between-group differences associated with these effects are sensitively captured when focusing on RT.

### 3.2 Age-related decline in early mnemonic stages

RTs to simple frequency changes (STEP) collapsed across d1, d8 and m6 showed substantial inter-individual variability and were also generally slower in the OLD than YOUNG group (Fig. 2C; W = 2766.5, p = .007, CI [15.9 92.9], mean OLD: 482 ± 115, YOUNG: 434 ± 108 ms). For each subject, the RT to the pattern emergence (RANREGn and RANREGr) was corrected by the RT to the STEP (median per session). This was done to control for individual variability in RT to a simple stimulus change, and thus isolate the computation time required to detect an emerging pattern.

We first analysed responses to REGn, as a measure of early mnemonic stages. We computed the median RT to REGn, collapsed across all sessions (Fig. 2D). The OLD group took longer than the YOUNG to detect the REGn patterns (RTs to REG: W = 2654.5, p = .03, CI [5.49 106.25]; mean OLD: 1807 ± .152; YOUNG: 1747 ± 137 ms), suggesting age-related decline of early mnemonic components (echoic/short-term memory) supporting pattern detection.

### 3.3 Age-related decline in long-term memory formation

Fig. 2E shows the median RT to REGn vs REGr for each block and session. We first focused on day 1 (memory formation stage). A mixed ANOVA with factors condition (RANREGn / RANREGr), block (block 1-3 of d1) and between-subjects factor group (OLD/YOUNG) yielded a main effect of condition [F(1, 130) = 82.52, p < .001, η_p_^2^ =.39], a main effect of block [F(2, 260) = 6.87, p = .002, η_p_^2^ = .05] and an interaction of condition by block [F(2, 260) = 7.57, p = .001, η_p_^2^ = .06]. This confirms the general pattern previously observed for this task (Bianco et al., 2020a): whilst RT to REGn patterns remains stable across blocks (no significant difference between blocks), RT to REGr becomes progressively faster with repeated exposure (block 1 vs 2 p = .004; block 1 vs 3 p < .001).

A main effect of group [F(1, 130) = 8.71, p = .004, η_p_^2^ = .06] and interaction of condition with group [F(1, 130) = 4.74, p = .031, η_p_^2^ = .04] confirmed that, across the 3 blocks of d1, the older group were slower overall than the younger group in detecting the REGr patterns (RT to REGr: OLD, mean 1725 ± 182 ms, vs YOUNG, 1622 ± 164, [t(130) = 3.40, p < .001]). An effect of ageing on REGn RT did not quite reach significance when focusing on the first day only (RT to REGn: OLD, mean 1809 ± 173 ms, vs YOUNG, mean 1761 ± 138 ms, [t(130) = 1.75, p = .08]), possibly due to some noise in the YOUNG data in block 2. No three-way interaction was found [F(2, 260) = 0.76, p = .467, η_p_^2^ = .01].

### 3.4 Computational modelling indicates that reduced performance among older listeners can be explained by reduced echoic buffer weight and faster LTM decay

Using a memory-constrained variant of Prediction by Partial Matching (Harrison et al., 2020), a computational model was optimised to fit the observed data on day 1 over blocks 1 to 3 of the ApMEM task for both OLD and YOUNG groups. We use this model to provide a formal simulation of early memory encoding and long-term memory formation characterising differences between the groups.

Fig. 3A shows the simulated RTs and the parameters optimised to fit the data of the YOUNG group (RMSE = 56.85). The decay kernel that these parameters generate is illustrated in Fig. 3C (right plot). Qualitatively, these parameters show close correspondence to those obtained when simulating responses of young participants on the same task in (Bianco et al., 2020a).

The optimisation of the data of the OLD group first examined whether a single parameter change from the values obtained for YOUNG could explain the differences in RT. The individually optimised parameters and their fit are given in Fig. 3B. It should be noted that, in several cases, the change of a single parameter affects the characteristics of multiple memory phases. No optimisation of only a single parameter managed to adequately fit the observed data for the OLD group, with all models possessing both high RMSEs reflecting an inability to recreate the trajectories of REGn and REGr responses, as displayed in Fig. 3B (left plot). In particular, parameters affecting the buffer, STM, or overall prediction noise, were unable to reproduce the decrease in learning rate exhibited in block 3 of the observed data for the REGr condition. While LTM weight was better able to account for this effect, it could not sufficiently increase simulated RTs for the REGn condition at the same time. Manipulating LTM half-life on its own was unable to produce a fit for either condition.

Next, pairs of parameters were optimised in turn to fit the data for the OLD group (a full list of parameter values and fit of models is given in supplementary material). Four parameter sets had better goodness of fit than the best-fitting single parameter optimisations when accounting for the increased number of parameters (i.e., with an AICc lower than that of LTM weight at 249.54). Each could plausibly fit the observed data, and each contained one parameter controlling properties of the buffer or short-term memory, and one controlling an aspect of long-term memory. The best-fitting combination (RMSE = 57.22; AICc = 247.26) was the optimisation of buffer weight and LTM half-life (0.93 and 232.86 s, respectively). The decay kernel described by these parameters diverges from that of the YOUNG group by having some down-weighting of memories formed within the buffer and more rapid long-term decay, as shown in Fig. 3C (right). These differences, and those of the other low-RMSE models fitting the OLD data (buffer duration and LTM half-life; buffer duration and LTM weight; STM duration and LTM half-life), indicate that the older group possesses weaker memory formation in both the immediate and long-term mnemonic phases and that both deficits are required to explain the differences observed between the two groups.

### 3.5 Auditory memory retained for up to 6 months in both older and younger listeners

Fig. 4A displays the RT advantage between REGr and REGn across all experimental sessions. The data revealed that the difference in RTA observed at the end of d1 (and modelled above) persisted when probed at d8 and at m6.

**Figure 4.**
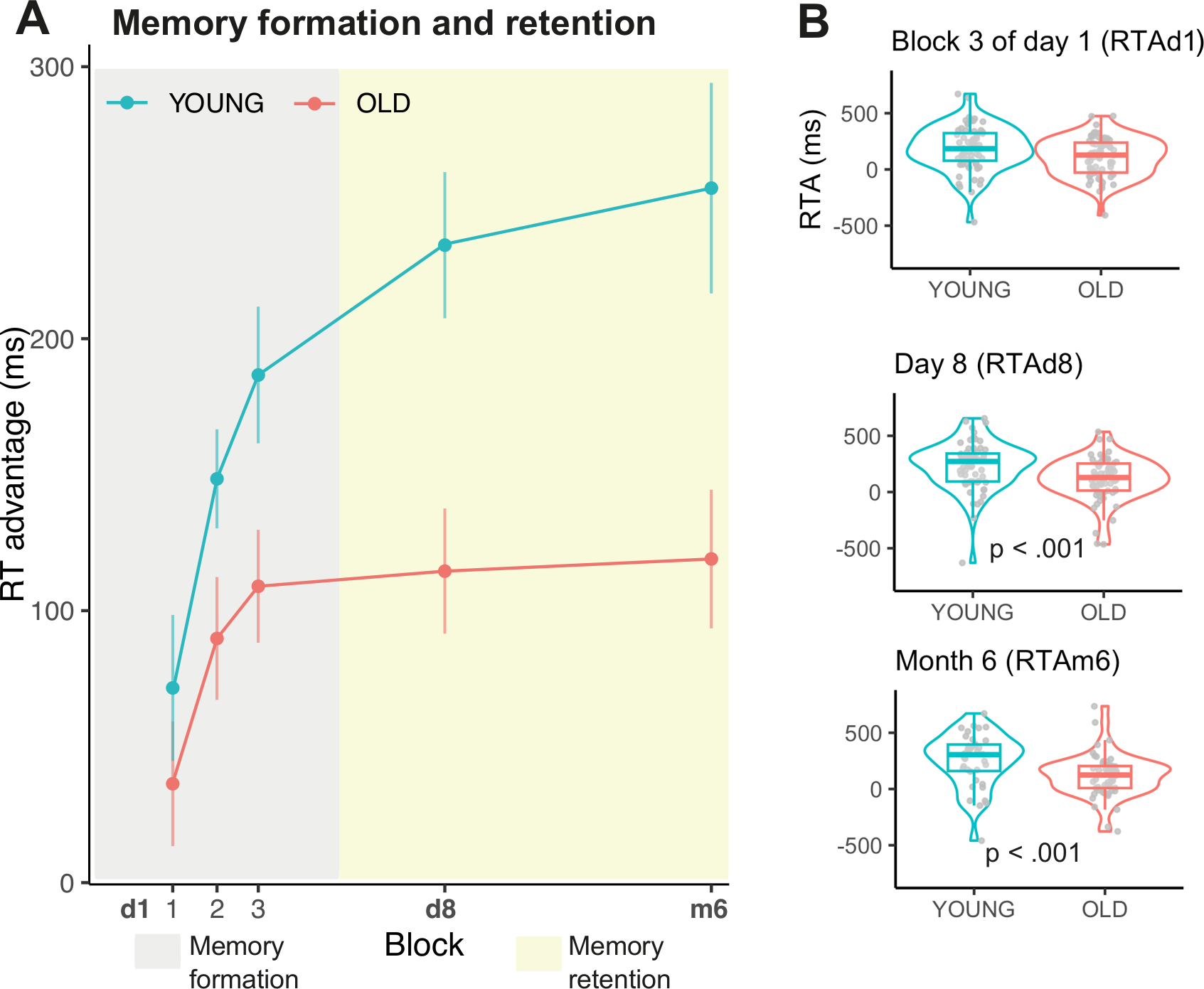
Auditory memory formation and retention in the young and older groups (ApMEM task). **(A) Long-term** memory dynamics across d1, d8 and m6 in young and older listeners, quantified as RT advantage (RTA) of REGr over REGn patterns. **(B)** Violin plots showing the individual data for the indexes extracted from the ApMEM task to quantify memory formation (the RTA computed over the last block of the first exposure on d1) and memory retention (the RTA computed on d8 and m6). Statistically significant differences between group means (Mann-Whitney U) are indicated.

To explicitly test for long-term memory retention, for each subject we compared the RTA at d8 and m6 to that observed in b1 (block 1 of d1, b1_d1). If listeners retained a lasting memory of REGr we expected RTA in d8 and m6 to be different from that in b1_d1. An ANOVA with factors block (b1_d1/ d8) and group yielded main effects of group [F(1, 130) = 8.69, p = .004, η_p_^2^= .06], block [F(1, 130) = 26.30, p < .001, η_p_^2^= .17], and no interaction [F(1, 130) = 3.24, p = .074, η_p_^2^= .02], indicating a greater RTA in d8 than in b1_d1 in both groups [t(131)= 5.03, p < .001; mean RTA OLD: b1_d1 36.4 ± 189, d8 115 ± .190 ms; YOUNG: b1_d1 71.6 ± 215, d8 234 ± 215 ms], and overall greater in the YOUNG than in the OLD group [t(130) = 2.94, p = .004].

An ANOVA with factors block (b1_d1 / m6) and group yielded main effects of group [F(1, 91) = 10.82, p = .001, η_p_^2^= .11], block [(1, 91) = 16.59, p < .001, η_p_^2^= .15], and no interaction [F(1, 91) = 1.52, p = .221, η_p_^2^= .02], indicating a greater RTA in m6 than in b1_d1 in both groups [t(92) = 3.90, p < .001; mean RTA OLD: b1_d1 30.2 ± 198, m6 119 ± 189 ms; YOUNG: b1_d1 89.5 ± 197, d8 255 ± 239 ms], and an overall greater RTA in the YOUNG group [t(91) = 3.28, p = .001]. This analysis shows that weaker LTM memory is formed in the OLD than in the YOUNG group, but both groups maintain non-decaying memories of the REGr patterns up to 6 months following initial memory formation.

Lastly, we compared the RTA in d1 block 3 (b3_d1), to those in d8 and m6 (Fig. 4B). A repeated measures ANOVA revealed a main effect of group only [F(1, 91) = 13.86, p < .001, η_p_^2^= .12], consistent with the overall larger RTA among the young listeners. There was no main effect of block, nor an interaction (block: [F(2, 182) = .97, p = .381, η_p_^2^ = .01]; interaction: [F(2, 182) = .93, p = .397, η_p_^2^= .01]), confirming a plateauing of the RTA after d1 in both groups – consistent with an enduring memory trace.

We also found that RTA in b3_d1 positively correlated with RTA in d8 (spearman’s rho = 0.198, p = 0.022) and that RTA in d8 correlated with RTA in m6 (spearman’s rho = 0.230, p = 0.026). This indicates good reliability of individual effects even in online settings.

Finally, we found no correlations between the age of the participants and the RTA in b3_d1, d8, or m6, in either the OLD (all p < .06), nor the YOUNG group (all p < .19). This may be due to the narrow age ranges used in the current study. However, the observed negative trend in the correlation between age and RTA in m6 in the older group (rho = −0.255, p = .06) suggests that future studies with larger sample sizes and a broader age range may potentially reveal critical age periods in implicit auditory memory decay.

To summarise, an effect of group (young vs. older) emerged across different time scales. In addition to weaker early mnemonic stages (Fig. 2D), the older group exhibited weaker long-term memory, as reflected by smaller RTA in b3_d1, d8 and m6. However, there was no evidence of a decline in memory e.g., between b3_d1 and d8 or d8 and m6.

### 3.6 No link between explicit and implicit auditory memory formation

Explicit memory for the REGr patterns was assessed at the end of d8 with a surprise familiarity task (Fig. 5A). Each REGr was presented once only amongst a large set of foils (REGn) and participants judged if the pattern was familiar. MCC (Matthews correlation coefficient) was used to measure the quality of subjects’ binary classification. Both groups exhibited above-chance performance (OLD: V = 1734, p < .001; YOUNG: V = 1765.5, p < .001), but the performance was poorer in older listeners (W = 1589, p = .040, CI [−.137 −5.16e-05], mean MCC OLD: .173 ±.217, YOUNG: .253 ± .183) (Fig. 5A). As in our previous findings (Bianco et al., 2020a), explicit memory scores did not correlate with the RTA observed in the last block of d1 (b3_d1; spearman’s Rho = −.025; p = .777) nor with that in d8 (spearman’s Rho = .056; p = .170). These analyses confirm the implicit nature of the RTA measures.

**Figure 5.**
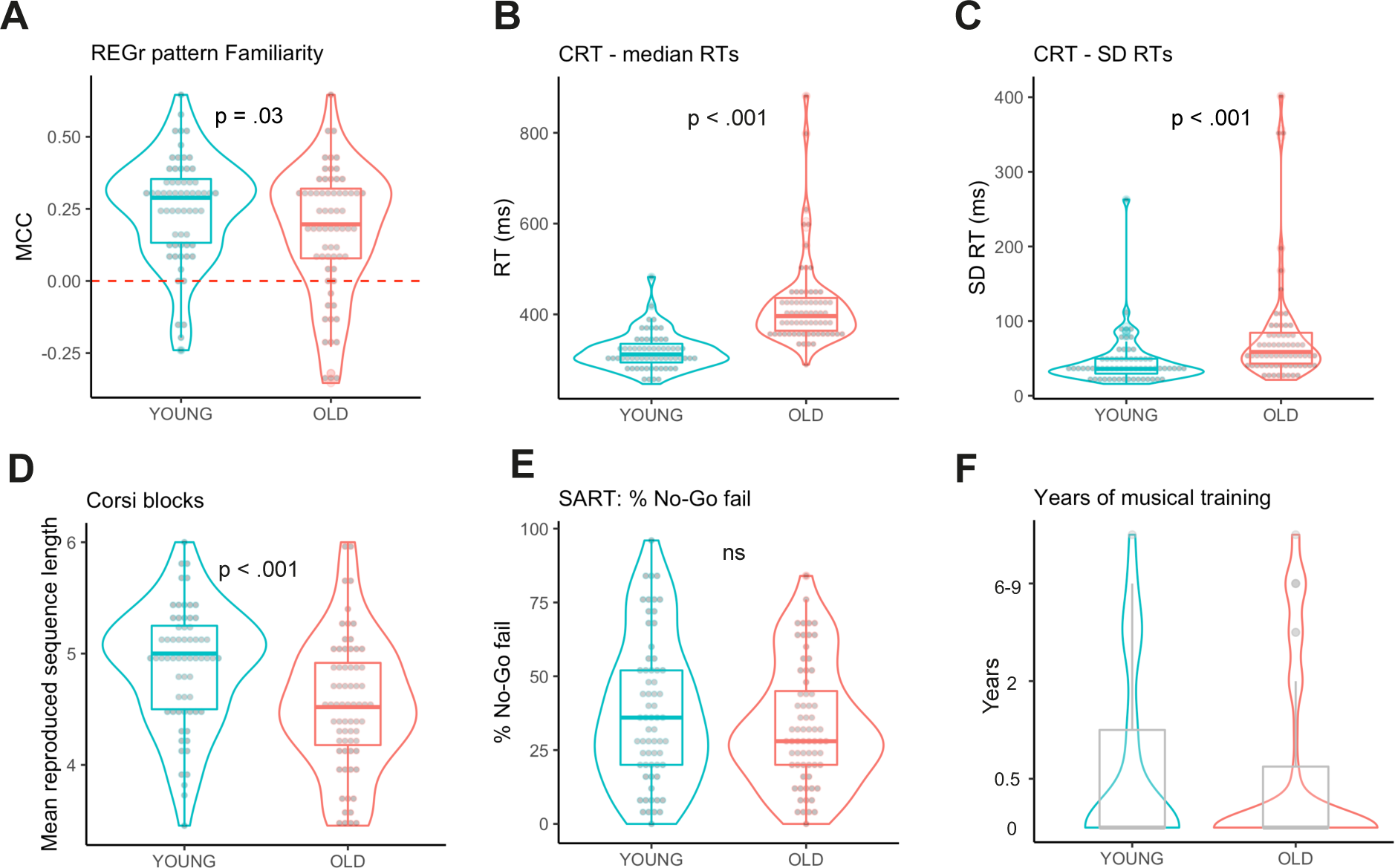
Distribution of performance, across the various tasks, in the young and older groups. Statistically significant differences between group means (Mann-Whitney U) are indicated. Worse performance in the OLD than in the YOUNG group was observed in **(A)** ApMEM explicit familiarity scores computed as Matthews correlation coefficient (MCC) on the reoccurring patterns and obtained at the end of day 8. The coefficient ranges between 1 (perfect classification) to −1 (total misclassification). **(B)** CRT both in terms of median and **(C)** variance of RTs, and **(D)** Corsi-blocks. Accuracy on the SART task **(E)** did not differ between groups. The two groups did not differ in terms of **(F)** musical training (W = 2076, p = 0.580).

### 3.7 Age-related decline in visual-sequence memory and processing speed, but no link with ApMEM RTA

At the group level, older participants showed slower median RTs in the CRT (W = 4000, p < .001, CI [68.80 100.09], mean OLD: 421 ± 97.2; YOUNG: 319 ± 40.5), and greater variability (SD of trials: W = 3227, p < .001, CI [11.63 27.06]) (Fig. 5B-C), reflecting a well-known effect of age-related processing speed impairment (Hultsch et al., 2002). The median RTs in CRT correlated with the RTs in our control STEP condition (Rho = .312, p < .001) confirming that STEP RTs are a good measure for correcting differences in baseline speed.

The older group exhibited worse performance in the Corsi blocks task (OLD vs YOUNG: t(130) = −3.56, p < .001, mean OLD: 4.54 ± .59; YOUNG: 4.80 ± .537), confirming the expected age-related decline in visual sequence memory (Fig. 5D) (Beigneux et al., 2007; Bianco and Chait, 2023; Fournet et al., 2012).

In line with previous findings on the SART (de Kerangal et al., 2021), there was no age-related decline in the sustained attention accuracy (% ‘no-go’ fail: W = 1967, p = .343, CI [−11.10 3.10], mean OLD: 33.5 ± 20.3; YOUNG: 38.1 ± 24.5) (Fig. 5E). As expected, RTs were slower in the OLD vs the YOUNG group in the ‘go-trials’ (W = 11073, p < .001, CI: [19.62 52.54], mean OLD: 378 ± 66.5; YOUNG: 348 ± 85.7). The speed-accuracy trade-off was similar between groups: accuracy in ‘no go’ trials was predicted by RT (χ^2^(1) = 25.29, p < .001), but not by group or the interaction between RTs and group (p > 0.1).

We conducted linear regression analyses to understand to what extent group-specific variance in ApMEM is predicted by performance on these cognitive tasks. We also included weekly hours of physical activity and years of musical training as possible predictors of ApMEM performance. Physical activity has been listed among the factors reducing the risk of cognitive and memory decline (Nyberg and Pudas, 2019; Sofi et al., 2011). Evidence from a meta-analysis has linked musical practice in healthy ageing with cognitive benefits both in domain-specific functions (auditory perception) and more general ones (Román-Caballero et al., 2018). For each outcome measure of ApMEM (Fig. 4B) and group, we performed a linear regression analysis with the predictors: CRT standard deviation, SART RTs, Corsi mean sequence length, and the above-mentioned demographic scores. None of the models was significant in the older group (all p-values > .11). A similar analysis in the younger cohort also yielded non-significant models (all p-values > 0.14). Overall, this pattern of results indicates that the variability in ApMEM is not driven by general processing speed, visuospatial sequential memory, or sustained attention, and might thus reflect age-related deficits specific to auditory memory.

## 4. Discussion

Despite its role in supporting fundamental aspects of auditory perception, how implicit sensory memory is affected by ageing remains poorly understood. Existing work has predominantly focused on verbal material and tasks that require cued reporting which are susceptible to factors such as rehearsal strategies or interference during long retention periods. For example, it is not known whether the “accelerated forgetting” effect recently characterised in older listeners (Elliott et al., 2014; Wearn et al., 2020) is specific to explicit memory or also extends to more basic auditory mnemonic representations.

Here, we introduce a paradigm that distils and quantifies age-related deficits in core memory mechanisms underlying auditory scene analysis. We compared younger (aged between 20 and 30 years old) and older adults (aged between 60 and 70 years old) with an RT-based implicit memory test. Participants are required to detect emerging regular patterns from random rapid sequences of tones. Patterns are novel in most trials (REGn), but unbeknownst to the participants, a few distinct patterns reoccur identically throughout the experimental sessions (REGr). The progressively growing RT advantage for REGr vs REGn demonstrates that mnemonic traces for the specific reoccurring patterns become more salient in memory through repeated presentation. The test is repeated 8 days and 6 months after initial exposure. Notably, the stimuli are arbitrary, and too fast to allow conscious tracking of the sequence events; thus, minimizing active tracking processes and mnemonic interference with real-world sounds.

We found that compared to young adults, older participants were slower in detecting novel patterns and exhibited a smaller RT advantage in detecting reoccurring patterns, indicating deficits in echoic/short-term memory and long-term memory formation. A computational model of auditory sequence memory fit to the data on the first day of exposure also suggested age-related limitations in both early and long-term mnemonic components. However, in contrast to mounting demonstrations of accelerated forgetting of verbal material with ageing, here older adults maintained stable memory traces for the reoccurring patterns – an unaltered RT advantage – up to 6 months after the first exposure. An absence of a correlation between the RT advantage and the active recall abilities measured with familiarity ratings confirmed the implicit nature of the RT effects.

### 4.1 Processing speed does not explain the reduced auditory memory effect in the older cohort

The between-group difference in the derived RT-based measures of memory cannot be explained by general reduced processing speed in the aged cohort (Salthouse, 1996; Zajac and Nettelbeck, 2018). This is supported by three arguments: First, RTs were corrected by RT to a simple stimulus change (STEP condition) interspersed in the main ApMEM task. This ensured that performance in pattern detection was controlled for inter-individual biological (e.g., subject’s general state of vigilance, or the time taken to perceive an auditory change, to generate a response) or equipment-based differences (e.g., keyboard latency) introducing non-memory specific variability. Second, the worse performance of older than younger adults in the control choice-RT task (CRT) showed the expected age effect on processing speed, and it correlated with the ApMEM STEP control condition, confirming the latter as a valid measure for correcting general differences in baseline speed. Finally, the regression analyses, including control tasks as predictors of ApMEM performance, showed that CRT did not contribute to variability in any of ApMEM-related memory measures. This, together with the absence of correlations between the ApMEM memory measures and tasks associated with sustained attention (SART) and visual-sequence memory suggest that between-group differences in ApMEM reflects age-related deficits specific to auditory memory.

Online RT measurements are subject to limitations due to the variability in the participants’ equipment (Anwyl-Irvine et al., 2020; Bridges et al., 2020) and engagement (Bianco et al., 2021). Although we implemented strict equipment requirements (see methods), residual components of equipment-driven RT delay might still contribute to the RT variance when measured online. Importantly, this variance is likely to be equally distributed across the groups (see methods and table S2).

### 4.2 Deficits in both early and long-term mnemonic stages contribute to age-related performance decline

Our measure of memory was RT which can be related to the computations inherent to dynamic sequence processing, and hence to the model output. Whilst accuracy measures were at ceiling by task design, older compared with younger participants exhibited overall slower RTs in response to novel patterns and, over time, formed a smaller RTA for the reoccurring patterns. Slower RTs in the older group could reflect a combination of slower access to stored information and less confidence due to noisier memory encoding. Whilst these factors could be reflected in both RT to REGr and REGn, the difference between the two conditions (the RTA) is likely to specifically reflect long-term memory representations. Modelling demonstrated that optimisation of parameters associated with both early and later mnemonic stages was necessary to accurately model older listeners’ performance.

We modelled the effect of ageing on memory by optimising the model memory decay kernel pertaining to: (1) a high-fidelity echoic memory buffer; (2) an STM phase; and (3) an exponentially decaying LTM phase. Each of these is associated with parameters describing their duration, relative weight, and rate of decay. Amongst the four best-fitting models, each contained one parameter controlling properties of the buffer or short-term memory, and one controlling an aspect of long-term memory. The best fitting model suggested that weaker echoic buffer weight (YOUNG: 1; OLD: .93), as well as more rapid long-term decay (YOUNG: 502.96 s; OLD: 232.86 s), contributed to the age-related performance decline on day 1.

The first ‘pre-perceptual’ stage of temporarily holding auditory information allows listeners to bind incoming events with the just heard ones to perceive a coherent representation of sequential sounds (e.g., a sequence of single tones as a motive). The computational modelling results support the interpretation that the overall slower detection of REGn patterns in older adults is a consequence of limited buffer / short-term memory components. This is in line with previous hypotheses (Herrmann et al., 2022) and consistent with neurophysiological evidence showing a somewhat weaker ability to represent information in echoic memory in older than young adults (Cooper et al., 2006; Pekkonen et al., 1996).

Parameters affecting the early stages of auditory memory alone could reproduce the age effect on responses to novel patterns but were unable to reproduce the diminished RTA for reoccurring patterns. A combination of reduced memory buffer and more rapid long-term decay (LTM half-life) best accounted for both aspects of performance differences between groups.

The decay kernel optimised for older adults’ data provides an initial model of how limitations at multiple stages of memory may explain different cognitive performance between populations. We focused on PPM-decay as the primary model for benchmarking auditory sequence tracking and learning. This model has successfully explained behavioural and neural responses to stimuli like those used here (Barascud et al., 2016; Bianco et al., 2020a; Harrison et al., 2020). Notably, relative to neural network (Huang et al., 2018) and Gestalt models (Narmour, 1990), PPM excels in simulating brain responses to more complex signals, such as music (Kern et al., 2022). Moreover, unlike alternative models (e.g., Skerritt-Davis and Elhilali, 2021), to the best of our knowledge, PPM-decay is the only model that has explicit decay parameters. In particular, the long-term decay seems to be necessary to simulate listeners’ performance on different auditory tasks (Bianco et al., 2020a; Harrison et al., 2020). The PPM model also provides some inherent reproduction of interference effects which are thought to co-exist with decay (Hardt et al., 2013; Lewandowsky et al., 2009).

The parameters for this model were optimised to fit blocks 1 to 3 of the ApMEM task performed on d1. As the constant long-term decay of the model predicts that memory should eventually reduce to zero, d8 and m6 are beyond the scope of this modelling in its current form. Modelling such time spans, while still being able to recreate effects within the first three blocks, would require a non-trivial addition that could account for memory consolidation over the intervening time periods.

Could the observed age-related memory effects arise from poorer hearing sensitivity in older listeners? We consider this unlikely for several reasons: firstly, the sensitivity to the presence of regularities was high among older adults and did not differ between older and young listeners. Secondly, during the instructions stage, participants were encouraged to adjust the sound volume to as high a level as needed, and the tone frequency range of 220-2000 Hz was intentionally chosen to be less affected by age-related hearing loss. This further guaranteed adequate audibility for all sounds. Finally, only participants who passed the headphone test (see methods) were included.

### 4.3 No evidence of long-term forgetting with ageing: memory traces to arbitrary tone patterns are retained for up to 6 months from initial exposure

The older compared with the young group formed weaker memory during the first day of exposure as quantified by a smaller RTA. However, just as observed in the younger group, the RTA in older listeners persisted for 8 days and 6 months after the initial exposure. This very long-lasting effect observed in both groups is noteworthy considering that the RTA was not driven by the explicit familiarity judgments obtained 6 months earlier.

There is growing interest in tests taxing memory circuit functionality at delayed recall because memory problems at this stage could indicate incipient dementia (Ryan and Frankland, 2022; Weston et al., 2018). That auditory patterns were not forgotten at delays of 8 days nor 6 months in the older cohort is in contrast to the body of work on accelerated long-term forgetting (ALF) for verbal material in ageing (Davis et al., 2003; Elliott et al., 2014; Manes et al., 2008; Mary et al., 2013; Wearn et al., 2020). ALF of verbal material has been reported also in pre-symptomatic autosomal dominant Alzheimer’s disease (Weston et al., 2018), patients with temporal lobe epilepsy (Blake et al., 2000), and it may represent a failure of memory consolidation processes (Hoefeijzers et al., 2013) due to altered integrity of hippocampal-neocortical (temporal) connections (Alvarez and Squire, 1994). One explanation for the discrepancy between verbal tasks and ApMEM may reside in the very low probability, compared to verbal material, that subjects encountered ApMEM-like sequences outside of the experimental sessions. This might have minimised phenomena such as forgetting due to interference with real-world stimuli (Davis and Zhong, 2017), which perhaps affects older more than younger adults (Wais and Gazzaley, 2014).

An alternative explanation can be found in the dichotomy between the active and implicit nature of the memorization processes inherent in verbal memory tasks versus the ApMEM task. While the former requires participants’ active engagement in memorization and subsequent recall, the latter relies on implicit memorization achieved through repetitive exposure. Notably, the memory effects observed in the ApMEM task, as quantified by RTA, did not correlate with the active recall of the REGr sequences in the familiarity task. This observation is in line with demonstrations in the visuomotor domain that implicit learning through repetition leads to memory retention for a remarkably long time (Kobor et al., 2017; Nemeth and Janacsek, 2011; Tóth-Fábera et al., 2023). These findings collectively suggest that implicit memory is rooted in robust biological substrates (Bailey et al., 2004; Ohno et al., 2011) less vulnerable to the availability of processing resources, attention, or interference, and so more preserved by ageing.

One important open question is why such long-term auditory implicit memory is overall resilient to time decay even later in life. The remarkable examples of preserved auditory memory for music in severe cases of dementia (Baird and Samson, 2009; Benhamou et al., 2021; Jacobsen et al., 2015) suggest that implicit auditory memory has a privileged status in the brain. In young listeners, implicit auditory memory based on repeated exposure has been demonstrated for many sound types, ranging from white noise (Agus et al., 2010; Dauer et al., 2022; Ringer et al., 2022), click trains (Kang et al., 2017), discrete sequences of tones (Bianco et al., 2020a; Bonetti et al., 2022; Herrmann et al., 2021; Leek and Watson, 1988), tone clouds (Agus and Pressnitzer, 2021), and naturalistic textures (Ringer et al., 2022; Woods and Mcdermott, 2018). All these studies capitalise on Hebb-type learning tasks (Hebb, 1961; Reber, 1989), whereby regardless of subject awareness, recognition of reoccurring patterns improves compared to novel ones simply due to reinforcement through repetition. Repetition is perhaps the simplest cue-inducing learning because it indicates the presence of patterns potentially relevant to behaviour (McDermott et al., 2011). Patterns have often a communicative function and are indeed implicitly learned through repetition in humans (Aslin, 2017; Saffran et al., 1996; Smalle et al., 2018) and non-human animals (Cazala et al., 2019; Hauser et al., 2001; Lu and Vicario, 2014; Soyman and Vicario, 2019; Wilson et al., 2013). The long-term memory of tone-sequences observed here even in older adults might thus reflect this primordial predisposition of the brain to remember patterns even when they sparsely reoccur.

## 5. Conclusions

Presenting a multi-stage auditory memory test that is scalable and easy to administer, we propose a paradigm shift in memory research beyond traditional verbal memory tests. We demonstrated that ageing is associated with poorer auditory echoic/short-term memory and long-term memory formation than young listeners, but not with forgetting. We speculate that ageing might affect frontal-auditory and hippocampal circuits underlying memory formation, but once formed auditory memories of rapid tone patterns remain accessible for months after the initial exposure even in older listeners. This result might be explained by the absence of interference with memory traces of arbitrary stimuli, unlikely to be encountered in daily life (Hardt et al., 2013; Ryan and Frankland, 2022), and suggests preserved long-term implicit auditory memory in ageing. Future studies combining human neuroimaging, animal models and synaptic simulations should shed light on the underlying circuits and neuronal mechanisms (Gershman, 2022; Ohno et al., 2011; Poeppel and Idsardi, 2022). Expanding the work to include a wider age range, from children to the elderly, could offer valuable insights into the currently understudied development and decline of auditory memory.

## Data availability

The datasets for this study are shared via https://osf.io/4d5se/?view_only=aa13d54c3ec248489bde41b9b562f470 and will be made publicly available upon publication.

## Authors contribution

RB: Conceptualization; Data curation; Formal analysis; Funding acquisition; Investigation; Methodology; Validation; Visualization; Roles/Writing - original draft; Writing - review & editing.

ETRH: Formal analysis; Methodology; Software; Validation; Visualization; Roles/Writing - original draft; Writing - review & editing.

MTP: Methodology; Resources; Software; Supervision; Validation; Visualization; Roles/Writing - original draft; Writing - review & editing.

MC: Conceptualization; Funding acquisition; Methodology; Project administration; Resources; Supervision; Validation; Visualization; Roles/Writing - original draft; Writing - review & editing.

## Supplementary material

**Table S1.**
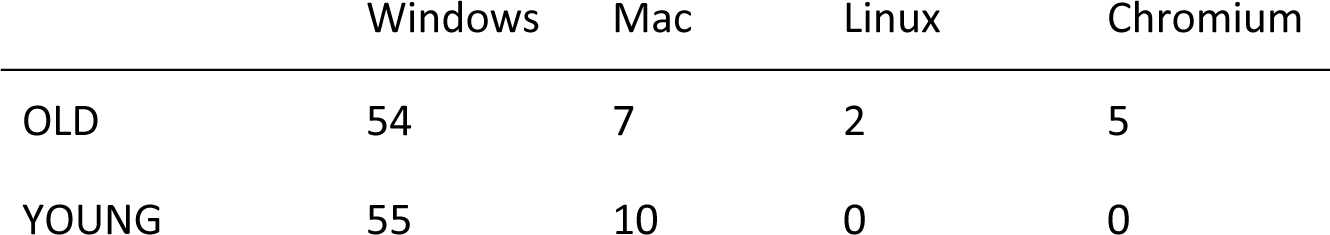
Counts of participants in the OLD and YOUNG groups per OS.

|  | Windows | Mac | Linux | Chromium |
| --- | --- | --- | --- | --- |
| OLD | 54 | 7 | 2 | 5 |
| YOUNG | 55 | 10 | 0 | 0 |

**Table S2.** Optimising parameter pairs to OLD ApMEM blocks 1 to 3.

| Parameter 1 | Value 1 | Parameter 2 | Value 2 | RMSE | AICc |
| --- | --- | --- | --- | --- | --- |
| Buffer weight | 0.86 | Buffer duration | 0.97 | 64.76 | 254.69 |
| Buffer weight | 0.90 | STM duration | 14.90 | 63.20 | 253.22 |
| Buffer weight | 1.00 | LTM weight | 0.006 | 61.34 | 251.43 |
| **Buffer weight** | **0.93** | **LTM half-life** | **232.86** | **57.22** | **247.26** |
| Buffer weight | 1.00 | Noise | 1.43 | 62.08 | 252.15 |
| Buffer duration | 0.80 | STM duration | 7.40 | 64.70 | 254.63 |
| **Buffer duration** | **0.69** | **LTM weight** | **0.009** | **58.52** | **248.60** |
| **Buffer duration** | **0.54** | **LTM half-life** | **185.94** | **57.53** | **247.59** |
| Buffer duration | 1.01 | Noise | 1.47 | 62.70 | 252.75 |
| STM duration | 13.07 | LTM weight | 0.005 | 61.67 | 251.75 |
| **STM duration** | **8.10** | **LTM half-life** | **217.05** | **58.62** | **248.71** |
| STM duration | 32.42 | Noise | 1.49 | 61.28 | 251.37 |
| LTM weight | 0.005 | LTM half-life | 544.37 | 61.63 | 251.72 |
| LTM weight | 0.005 | Noise | 1.31 | 61.72 | 251.80 |
| LTM half-life | 237.87 | Noise | 1.40 | 59.51 | 249.62 |

Values of parameters and their RMSE and AICc fit to RTs of the OLD group, when optimising pairs of parameters in turn to those of YOUNG. Multiple pairs of parameters produced plausible possibilities for differences between groups, with the best fitting parameters being buffer weight and LTM half-life. The best-fitting combinations (buffer weight and LTM half-life; buffer duration and LTM half-life; buffer duration and LTM weight; STM duration and LTM half-life) each contained one parameter controlling properties of buffer or short-term memory, and one controlling an aspect of long-term memory.

### Comparison of tone and pitch interval modelling

The simulation of participants’ responses reported in this paper modelled sequences using the absolute pitches of their tones. However, these sequences may also be represented by the size and direction of intervals between tones. While there are many situations where the intervalic content of an auditory sequence may be the more salient feature (Saffan et al., 2004) (*e.g.,* in more complex stimuli, such as music, where relative pitch may be an advantage in pattern recognition), the nature of the RANREG stimuli used in the present experiment results in similar outputs when using either representation of stimuli sequences.

In the RANREG stimuli used in this experiment, repetition of patterns in interval occur concurrently with repetitions in absolute tone pitch. When a REG pattern is established, it repeats both its tone and interval information exactly; RAN sections contain both random pitch and interval content. These stimuli do not contain the common scenarios in which a distinction between absolute pitch and interval provides different information, such as the transposition of a pattern’s pitches but maintaining interval sizes. As a result, simulated RTs using absolute tone values and intervals are highly correlated and produce a similar fit to the task data. YOUNG: *r*(268) = 0.86, *p* < .001; OLD: *r*(268) = 0.86, *p* < .001.

**Figure S1.**
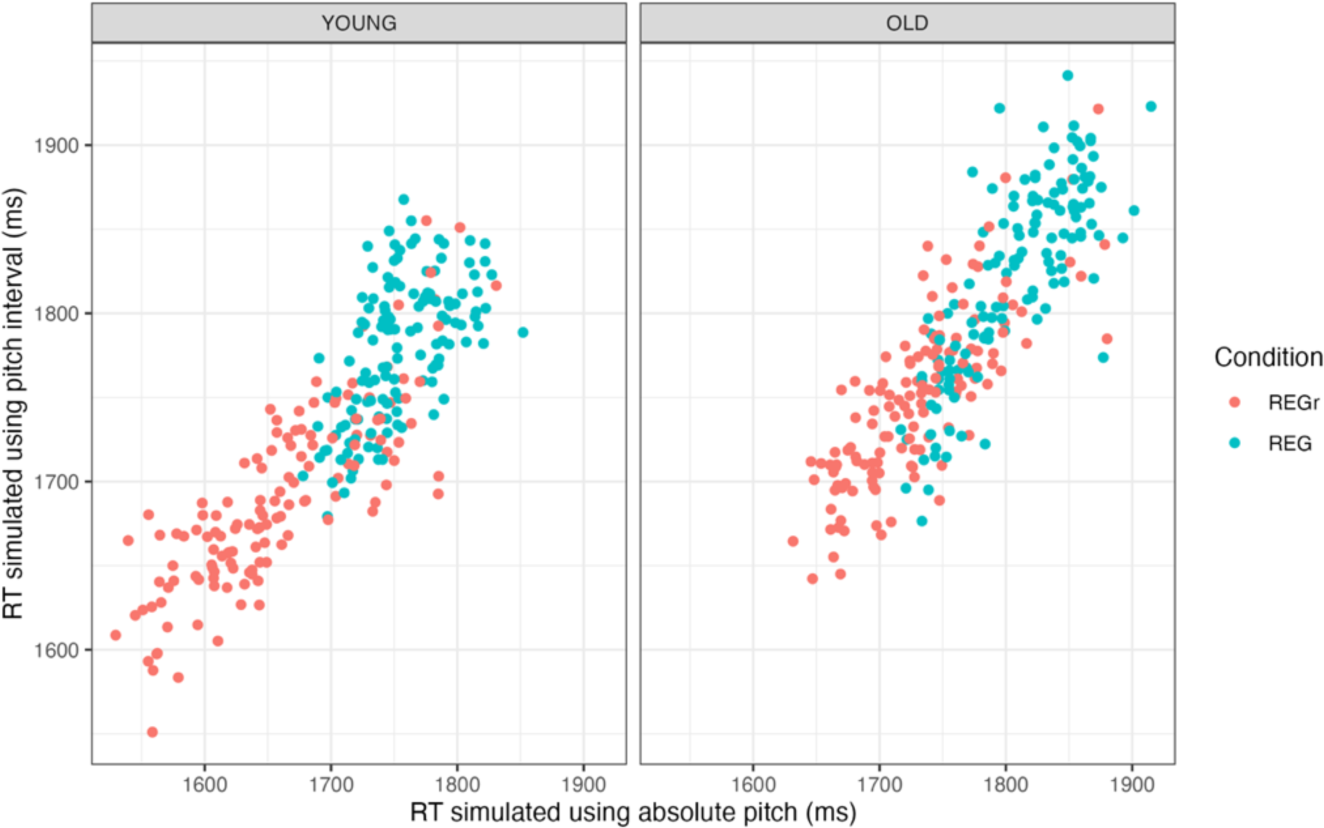
Comparison of simulated RTs using representations of absolute tone values and tone intervals. Modelling for YOUNG and OLD groups used the parameters obtained through optimisation to observed RTs when using the tone representation (see Figure 3). Parameters of the OLD group differ from those obtained for the YOUNG in *buffer weight* and *LTM half-life*.

**Figure S2.**
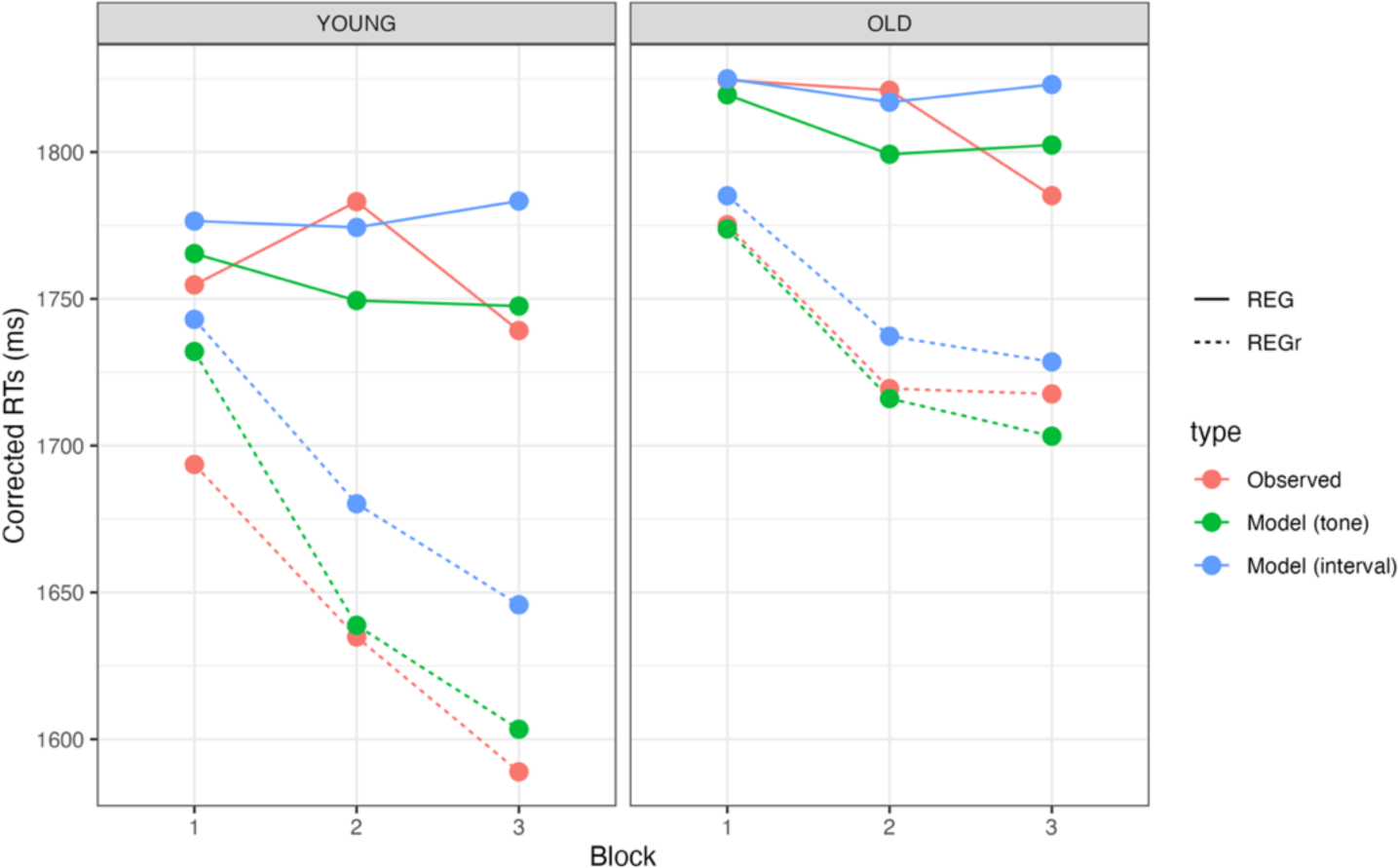
Observed and simulated RTs using representations of absolute tone values and tone intervals. Modelling for YOUNG and OLD groups used the parameters obtained through optimisation to observed RTs when using the tone representation.

